# Physical traits of supercompetitors in cell competition

**DOI:** 10.1101/2024.10.14.618306

**Authors:** Logan C. Carpenter, Shiladitya Banerjee

**Affiliations:** Department of Physics, Carnegie Mellon University, Pittsburgh, PA, USA

## Abstract

Cell competition is a fitness control mechanism in tissues, where less fit cells are eliminated to maintain tissue homeostasis. Two primary mechanisms of cell elimination have been identified in cell competition studies: contact-dependent cell death and mechanical compression-driven apoptosis. While both occur in tissues, their combined impact on population dynamics is unclear. Here we develop a cell-based computational model to study competition between two cell types with differing physical properties. The model integrates cellular mechanics with cell-cycle dynamics, contact-induced apoptosis, and cell extrusion via mechanical stress. Using this model, we explored how differences in physical traits between cell types influence competitive interactions. Our findings show that differences in cell compressibility alone can drive mechanical competition, with stiffer cells outcompeting softer ones in otherwise identical populations. Surprisingly, mutations that reduce cell stiffness, combined with decreased contact inhibition of proliferation, can create a “soft” super-competitive mutant. We demonstrate that changes in apoptosis sensitivity, cell adhesion, and cell size significantly affect growth potential and susceptibility to apoptosis. Furthermore, mutant cell colonies require a critical colony size, dependent on cell compressibility, to overtake the surrounding wild-type tissue. For colonies below the critical size, the elimination process is stochastic, driven by a protrusive finger-like instability in the interface between two cells that promote invasion of the supercompetitors.

## I. INTRODUCTION

Cell competition is a quality control process in tissues, where less fit cells are eliminated by their fitter neighbors [1– 3]. This phenomenon plays a key role in tissue development, maintenance of tissue homeostasis, and in the progression of cancer [4]. First discovered in *Drosophila* [5], cell competition was initially thought to act as a homeostatic mechanism, removing mutant cells with lower proliferative fitness. Later studies revealed mutants with a competitive advantage over wild-type cells that drive the elimination of wild-type cells [6]. A critical question is: what are the biophysical traits of mutations that provide a competitive advantage or disadvantage over normal cells?

Two primary modes of cell competition have been identified in previous studies: mechanical competition that can act over a distance [7–10], and contact-based competition, which directly affects the fate of neighboring cells through intercellular signaling [1, 11–14]. Mechanical competition is typically driven by differences in growth rates between “winner” and “loser” cell types, creating mechanical stress on the losers, ultimately leading to their apoptosis or extrusion [7, 15]. However, additional cellular factors such as cytoskeletal rigidity, membrane tension, preferred cell density, and cell-cell adhesion can also influence a cell’s ability to resist mechanical forces. A related determining factor in mechanical competition is contact inhibition of proliferation [16], which describes how cell growth and proliferation are inhibited in crowded environments [17–21]. Inducing mutations that downregulates the cellular contact inhibition have been shown to produce oncogenic-like supercompetitors [22– 24]. While mechanical and contact-based elimination processes often co-exist in tissues [25], their relative contributions to population size and cell survival are not well understood. This presents an experimental challenge, given current limitations in isolating the diverse biophysical factors, and in connecting single-cell behaviors to population-level dynamics [26]. To address this, theoretical models can provide valuable insights into how cell-level properties and processes of proliferation and death influence tissue-scale outcomes in cell competition.

Previous theoretical studies on cell competition have considered predator-prey interactions in continuous models [27, 28], the impact of mechanical stress on cell growth and apoptosis in continuum models [7, 15, 29], and cell-based models using the framework of vertex and Cellular Potts models [30– 33]. Cell-based models are particularly useful for linking single-cell behaviors to population-level dynamics [26]. Existing models have examined how factors like cell mechanics [31, 34], cell-cell adhesion [31, 35], contractility [36], and death signaling [32] influence the outcomes of cell competition betwewn “winner” and “loser” phenotypes. However, these models have not addressed the combined effects of mechanical and contact-based apoptosis on population survival. We address this gap by integrating mechanical and biochemical modes of competition at the cellular level, allowing us to identify the physical cell properties that predict the emergence of supercompetitors.

We advance our recently developed cell-based model for epithelial tissue growth and homeostasis [37] to now simulate multiple cell types and their elimination via contact-based and mechanical apoptosis. Our model comprises two integrated layers: one that captures the physical dynamics of tissues and another that determines cellular fate through probabilistic decision-making rules. The physical layer employs the framework of the cellular Potts model [38], while the decision-making layer establishes rules for cell-cycle regulation, contact inhibition of proliferation, and cell elimination via apoptosis and extrusion. Unlike previous cellular competition models, we do not assign winner or loser fates to cell types, but rather determine the cellular traits that lead to the emergence of losers or winners in cell competition. Our approach implements contact inhibition through crowding-induced suppression of cellular growth rates and considers the interplay between mechanical and contact-based cell competition. Additionally, we incorporate biphasic cell-cycle progression that allow for tunable growth and introduce distinct probabilities for apoptosis based on cell density and the proportion of heterotypic circumferential contact. Collectively, our model integrates feedback between mechanical pressure and the rates of cell growth, division, and apoptosis within the context of cell competition.

Using this model, we predicted key characteristics of supercompetitive cell colonies. These include softer cells with higher fitness, the advantage conferred by a faster cell-cycle and lower contact inhibition of proliferation, and the identification of a critical colony size necessary for invasion. The size-dependent elimination of fitter mutants underscores the crucial role of fitness monitoring within wild-type epithelial tissues. Essentially, the earlier the competitive mechanisms activate to eliminate a highly proliferative mutant, the better the chances for the wild-type tissue to prevail. Consequently, mutants with shorter cell-cycles have a higher likelihood of reaching the critical size. Interestingly, we discovered that mutant cells that are softer than their wild-type counterparts, can succeed by reducing their sensitivity to crowding. Taken together, our works reveals the intricate interplay between mechanical and decision-making elements of cells in regulating population growth, survival and extinction.

## II. CELL COMPETITION MODELING FRAMEWORK

To study the dynamics of cell competition on the scale of individual cells, we implemented a two-layered computational model: (1) A physical layer simulates the mechanics of the tissue via energy minimization, and (2) a decision-making layer that simulates cell-autonomous dynamics such as growth, mitosis and apoptosis (Fig. **1**). This framework allows us to characterize individual cell types by their expressed behaviors and physical characteristics informed by experiments [25].

**Fig. 1.**
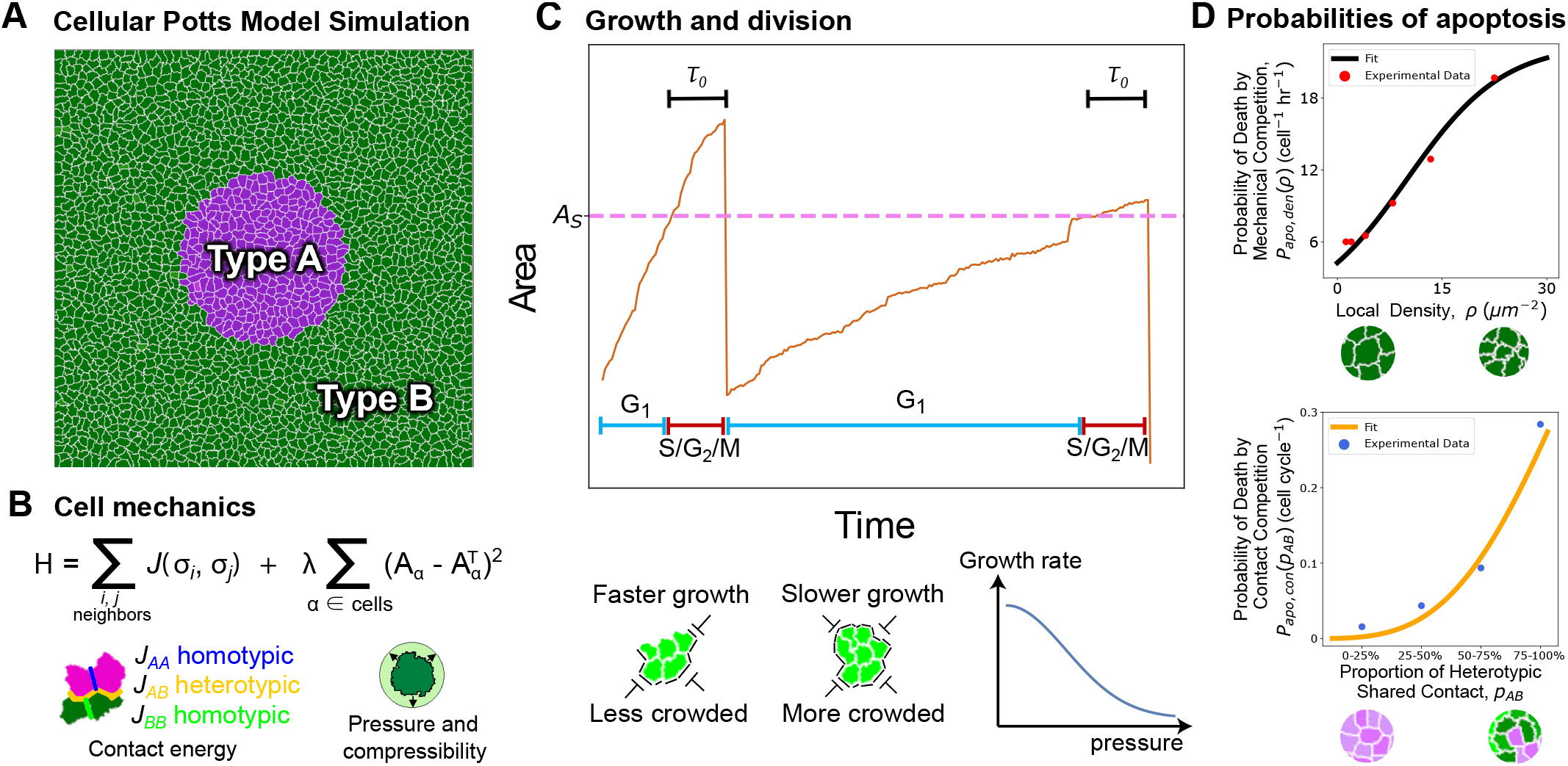
Overview of the computational model. (A) Snapshot at the beginning of a simulation. The purple cells, type A, represent the mutant population. The green cells, type B, represent the wild type population; throughout this study, the default parameters defining these cell populations remain constant except where changes to type A are explicitly stated. (B) The Hamiltonian used within the Cellular Potts model to determine how the system mechanics evolves. (left) Diagram of contact energies for heterotypic and homotypic contact. (right) Schematic depicting forces promoting growth due to mechanical pressure and compressibility *λ* of cells. (C) Rules for growth and mitosis. (upper) A plot representing a cell’s area trajectory over two cell cycles. The cell remains in the G1 phase until its area exceeds the sizer threshold at which point it transitions to the S/G2/M phase for a duration equal to the timer threshold, *τ*_0_. (lower) The diagrams depict (left) cells with space to expand into growing faster and (right) cells with no space to grow into growing slower. The qualitative effects of these two scenarios for crowding is represented in the above area plot. Growth dynamics are determined by changing the target area of the cell, *A*^*T*^, at a rate that decreases with increasing pressure on the cell. (D) Best fit curves for the probability of apoptosis as a function of (upper) local density (Eq. 5) (data from Ref. [31]) and (lower) the proportion of heterotypic shared contact (Eq. 6) (data from Ref. [12]).

### A. Physical Dynamics

The physical layer of the model is simulated using the Cellular Potts Model (CPM) [38], a computational framework widely used in the study of cell behaviors and tissue morphogenesis [39] (Appendix A). The CPM simulates a cell on a discrete lattice as a collection of lattice-sites assigned a unique positive integer identifier *n*_*id*_, with *n*_*id*_ = 0 used for the extracellular medium (Fig. **1**A). The mechanical energy of the system is given by

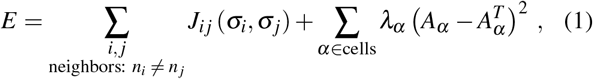

where the first sum captures the contact energy arising from cell-cell or cell-medium adhesion, and the second term describes the area elastic energy (Fig. **1**B). The cell type at lattice site *i* is given by *σ*_*i*_, which can take on values *σ*_*i*_ = { *A* : type A, *B* : type B, *M* : Medium}(Fig. **1**)A). Then the contact energy between lattice-sites *i* and *j* can take one of five possible values *J*_*i j*_(*A, A*) = *J*_*AA*_, *J*_*i j*_(*A, B*) = *J*_*AB*_, *J*_*i j*_(*B, B*) = *J*_*BB*_, *J*_*i j*_(*A, M*) = *J*_*AM*_, or *J*_*i j*_(*B, M*) = *J*_*BM*_. Due to energy minimization, a lower value of contact energy implies a greater adhesion strength. As a default, we assumed the greatest homotypic adhesion between type B cells (wild-type), and a lower homotypic adhesion between type A cells (mutants), and the least adhesion between cells and the medium *J*_*BB*_ *< J*_*AB*_ *< J*_*AA*_ *< J*_*BM*_ = *J*_*AM*_. The second sum in Eq. (1) iterates over each cell in the simulation to calculate their area elastic energies. For a given cell *α*, this is done by penalizing deviations of the true cell area *A*_*α*_ from its preferred (target) value 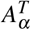, multiplied by the compressional modulus *λ*_*α*_ = { *λ*_*A*_ : *α* is type A, *λ*_*B*_ : *α* is type B}. Both adhesion and elastic energy parameters are varied in simulations.

The physical layer progresses in time by minimizing the mechanical energy *E* via implementation of the Metropolis-Hastings algorithm [40, 41]. This is done by a semi-stochastic flipping of the cell identity at individual lattice sites; for instance, the algorithm considers adjacent sites *i* and *j* (with distinct cell identities *n*_*i*_ ≠ *n* _*j*_), and calculates the change in system energy resulting from the change of cell identity at *i* to 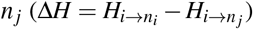. The probability of accepting the flip is given by

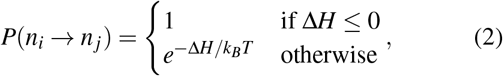

where *k*_*B*_*T* is the thermal energy scale of the Metropolis-Hastings algorithm.

### B. Decision-making rules

The physical layer of the CPM is coupled to a decision-making layer that implements various cell-autonomous rules between each simulation timetep, such as altering the cell’s target area, progressing the cell cycle phase, executing mitosis, initiating apoptosis, and performing extrusion of cells. For a given cell *α*, growth dynamics are governed by increasing the cell’s target area over time as [31, 37] (Fig. **1**C),

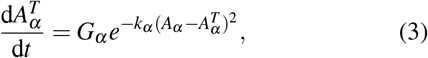

where *G*_*α*_ represents the uncrowded growth rate (or growth rate in isolation) of cell *α*, when 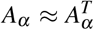, and *k* is a phenomenological parameter that captures cellular sensitivity to crowding or contact inhibition of proliferation. The *α* subscript on *G* and *k* is to distinguish the dependence of these values on cell type. When a cell lacks the physical space to grow, then the cellular pressure 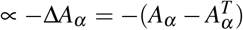 will increase, which in turn causes an exponential decay of the rate of growth in proportion to *k*. Furthermore, crowding leads to a decreased rate of division as the cell cycle progression from G1 to S/G2/M phase depends on cell size and growth rate.

We model cell cycle dynamics using a G1 sizer model [42, 43], where the cell cycle is split into two phases, G1 and S/G2/M (Fig. **1**C). A newborn cell begins to grow in the G1 phase and remains there until its area exceeds a threshold value, *A*_*S,α*_, where *α* distinguishes the value for a given cell type. Once *A*_*α*_ ≥ *A*_*S*_, a fixed countdown timer begins, which is the duration of the S/G2/M phase. After *τ*_0_ time has elapsed in the S/G2/M phase, the cell divides along the semimajor axis. The target areas of the newborn daughter cells are set to their actual area, 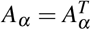. Both *τ*_0_ and *A*_*S,α*_ are drawn from Gaussian distributions that are derived from experimental data on Madin-Darby canine kidney (MDCK) cells [43]. Hence in our model, contact inhibition of proliferation results from tissue crowding halting cycle progression from G1 to S/G2/M [17].

We implement three different modes of cell elimination in our model, based on experimental observations: density-dependent apoptosis [25], contact-dependent apoptosis [12], and live-cell extrusion [18]. The initiation of density- and contact-dependent apoptosis are governed probabilistically by the local cellular environment, while extrusion operates deterministically, driven by the interplay between cellular and tissue-level states. We define the local density of a cell *α* as the sum of the inverse areas of the cells in direct physical contact with the cell *α*:

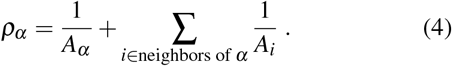

The functional form for the dependence of the probability of apoptosis on local cell density is obtained by by fitting a logistic curve to the experimental data of apopotosis probabilities obtained by Bove et al. [25] on MDCK cells (Fig. **1**D)

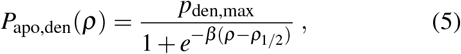

where *p*_den,max_ is the maximum probability of apoptosis, *ρ*_1*/*2_ is the cell density at half-maximum probability and *β* is the sensitivity to density-dependent apoptosis. Similarly, the probability of contact-dependant apoptosis is obtained by fitting a Hill function to the data obtained by Levayer et al. [12] in *ex vivo Drosophilia* wing discs (Fig. **1**D):

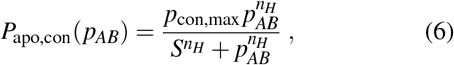

where *p*_*AB*_ is the proportion of cell perimeter in heterotypic contact, *p*_con,max_ is the maximum probability of apoptosis, *S* is the steepness parameter, and *n*_*H*_ is the Hill coefficient [31]. On initiation of apoptosis, regardless of the method, the cell’s target area is set to zero, which leads to a rapid decrease in area until eventually being removed from the tissue (see Appendix B for details). Lastly, a cell *α* is eliminated via extrusion when its area is less than a quarter of its cell type’s mean area at that time step, *A*_*α*_ < ⟨*A*⟩ _*α*_ /4. Upon meeting this condition, the cell is promptly deleted to mirror the short timescale of live cell extrusions, and its lattice sites are converted to medium lattice sites.

Stacking the physical and decision-making layers produces a predictive cell-autonomous model that can emulate the spatiotemporal tissue dynamics of cell competition. We now use this model to investigate the growth potential of a small colony of mutant cells surrounded by a tissue of wild-type cells.

## III. RESULTS

### A. Differential compressibility drives cell competition

A viable cell must sustain adequate rigidity to facilitate growth and division. Prior investigations have shown that MDCK cells, upon depletion of the polarity gene *Scribble*, exhibit reduced rigidity and are consequently outcompeted by wild-type MDCK cells [25, 31]. Furthermore, differential cell stiffness is also associated with mechanical competition between normal and bacterially-infected cells [44], and also in cancer cell invasion [45–48]. To test the predictive capability of our model, we investigated the emergence of disparities in cell fitness associated with differential compressibility. We simulated the growth of a small colony of Type A cells with a different compressibility compared to the surrounding Type B cells (Fig. **2**A, Videos 1-3). To isolate the effects of compressibility, all other cellular parameters for types A and B were kept fixed at their default values. See Tables I-II and Appendix C for details on parameter choices.

**TABLE I.**
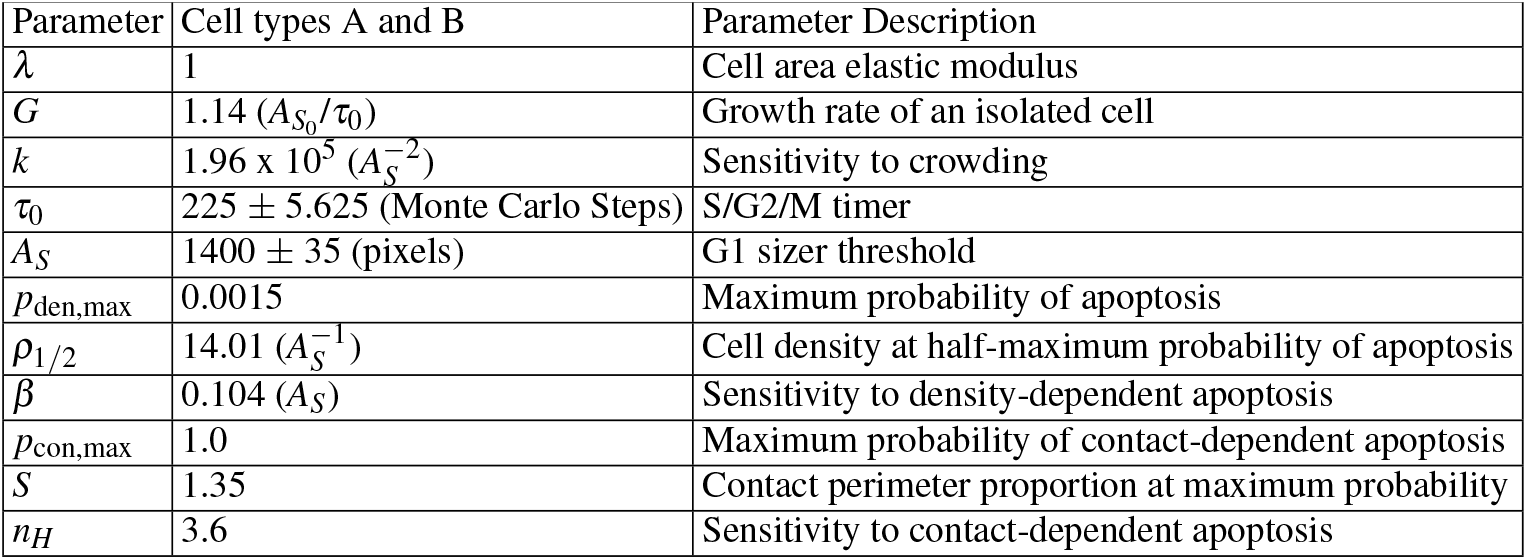
Default parameter values used in model simulations.

**TABLE II.** Default contact energies used in model simulations.

| Contact Energies | X = Type A | X = Type B |
| --- | --- | --- |
| $J_{XX}$ (Homotypic) | $2.39 \times 10^{-4} (\lambda A_S^{3/2})$ | $1.15 \times 10^{-4} (\lambda A_S^{3/2})$ |
| $J_{XY}$ (Heterotypic) | $1.91 \times 10^{-4} (\lambda A_S^{3/2})$ | $1.91 \times 10^{-4} (\lambda A_S^{3/2})$ |
| $J_{XM}$ (Cell-medium) | $2.86 \times 10^{-4} (\lambda A_S^{3/2})$ | $2.86 \times 10^{-4} (\lambda A_S^{3/2})$ |

**Fig. 2.**
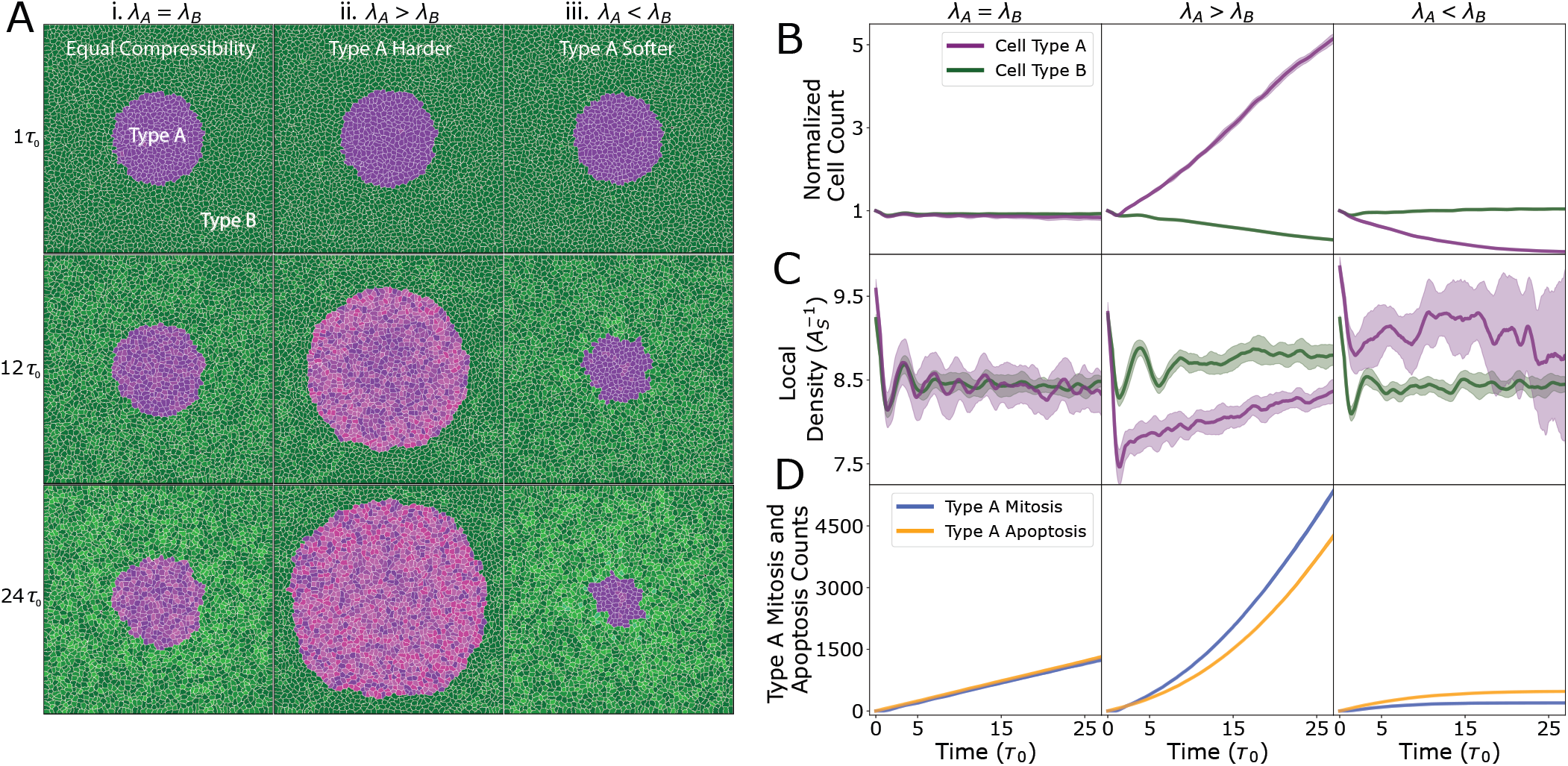
Differential compressibility alone drives cell competition. (A-D) Each column represents a different relative value of compressional elastic modulus of cell type A, *λ*_*A*_, with respect to the comparessional modulus of cell type B, *λ*_*B*_ (with *λ*_*B*_ = 1 for all simulations). All simulations start from the same initial configuration. (A) Snapshots taken of simulations at *t* = *τ*_0_, 12*τ*_0_, and 24*τ*_0_ (first, second, and third rows respectively). Columns indicate the compressional elastic modulus of type A cells when compared to type B cells: (i) *λ*_*A*_ = *λ*_*B*_, (ii) *λ*_*A*_ *> λ*_*B*_, (iii) *λ*_*A*_ < *λ*_*B*_. (B-D) The plots show the (N = 5) average value (darker line) and standard deviation (lighter color). (B) Normalized cell count (*N*(*t*)*/N*(0)) as a function of time (purple for type A and green for type B). The more compressible cells are eliminated from the tissue while the equally compressible cells survive (C) Local density as a function of time (purple for type A and green for type B). Cells with a lower relative elastic modulus are compressed above their homeostatic density, which increases their probability of mechanical apoptosis. (D) Counts of mitosis and apoptosis for type A cells. (right) With *λ*_*A*_ < *λ*_*B*_, type A cells are eliminated, in part, due to their low division count, which indicates an inability to grow. See tables I and II for a list of default model parameters.

When the system is composed of cells of identical compressional modulus (*λ*_*A*_ = *λ*_*B*_), the population dynamics of type A and type B cells are fairly identical across time (Fig. **2**B-i, Video 1), the average local densities are approximately similar (Fig. **2**C-i), and the rates of mitosis and apoptosis of the type A colony are nearly equal over time (Fig. **2**D-i). The simulation snapshots in Fig. **2**A-i show minor morphological changes in colony A, but qualitatively, colony A size is unchanged over a long timescales. In contrast, simulations with softer type B cells *λ*_*A*_ *> λ*_*B*_ (Fig. **2**A-ii, Video 2), which are compressed above their homeostatic density (Fig. **2**C, compare green curves in panel i with panel ii), are subsequently eliminated via mechanical competition (Fig. **2**B-ii). Consequently, we observe a higher rate of mitosis in Type A cells compared to the rate of apoptosis (Fig. **2**D-ii). In the case of softer type A cells *λ*_*A*_ *< λ*_*B*_ (Fig. **2**A-iii, Video 3), they are compressed beyond their equilibrium density (Fig. **2**C, compare purple curves in panels i and iii) and are then eliminated preferentially (Fig. **2**B-iii). This results from a higher rate of apoptosis of Type A cells compared to the rate of mitosis. These data collectively indicate that cell compressibility is a key factor in determining the viable mechanical pressure at which growth and proliferation occur, and when a cell’s environmental pressure exceeds this viable pressure, then growth effectively halts and the cell cycle is arrested in the G1-sizer phase. Since all other parameters are being held equal in these simulations, the higher density of cells with lower compressional modulus represents a decrease in fitness due to it also lowering the rate of division and increasing the rate of elimination: small cells cannot proliferate due to halting in the G1-sizer phase, and cells with a high local density have a greater probability of death (Eq. (5)).

### B. Trade-off between cell compressibility and sensitivity to crowding controls the outcome of cell competition

Our simulations thus far indicate that cells with a higher compressional elastic modulus emerge as supercompetitors, when all other cellular properties are identical. This finding contrasts with expectations for cancer cells, which tend to be softer than normal cells [46–48] but still outcompete them. We then explored the extent to which a reduction in fitness due to a lower compressional modulus could be counterbalanced by an increase in fitness driven by another parameter. Since cancer cells are characterized by a lower contact inhibition of proliferation [16, 49, 50], we varied the ratio of the sensitivity to crowding, *k*_*A*_*/k*_*B*_, alongwith the ratio of compressional moduli *λ*_*A*_*/λ*_*B*_, in order to investigate their combined effects on the fitness of the mutant colony A.

To characterize the fitness of colony A, we measured the net growth rate *α*, defined as d*N/*d*t* = *N*_0_ exp(*αt*), where *N*(*t*) is the cell count of type A cells at time *t*. We determined *α* by fitting an exponential function to the average (*n* = 6) population size (Fig. **3**). By varying *λ*_*A*_*/λ*_*B*_ and *k*_*A*_*/k*_*B*_, we mapped the fitness landscape (Fig. **3**), identifying regions where type A tissues exhibit super-competitive behavior (red points, *α >* 0), where type A tissues are eliminated (blue points, *α* < 0), and where type A and B tissues coexist in an unstable equilibrium (on dashed line, *α* ≈ 0). As we discussed in the previous section, an increase in compressional elastic modulus is associated with an increase in cellular fitness. However, we find that the decrease in fitness from softer (more compressible) cells can be compensated for by a proportional reduction in sensitivity to crowding, such that there is a net increase in fitness (Fig. **3**, purple shading). We thus identify a region in *λ* -*k* phase space that correspond to soft supercompetitors, where softer type-A cells are able to outcompete harder type-B cells due to their lower sensitivity to crowding.

**Fig. 3.**
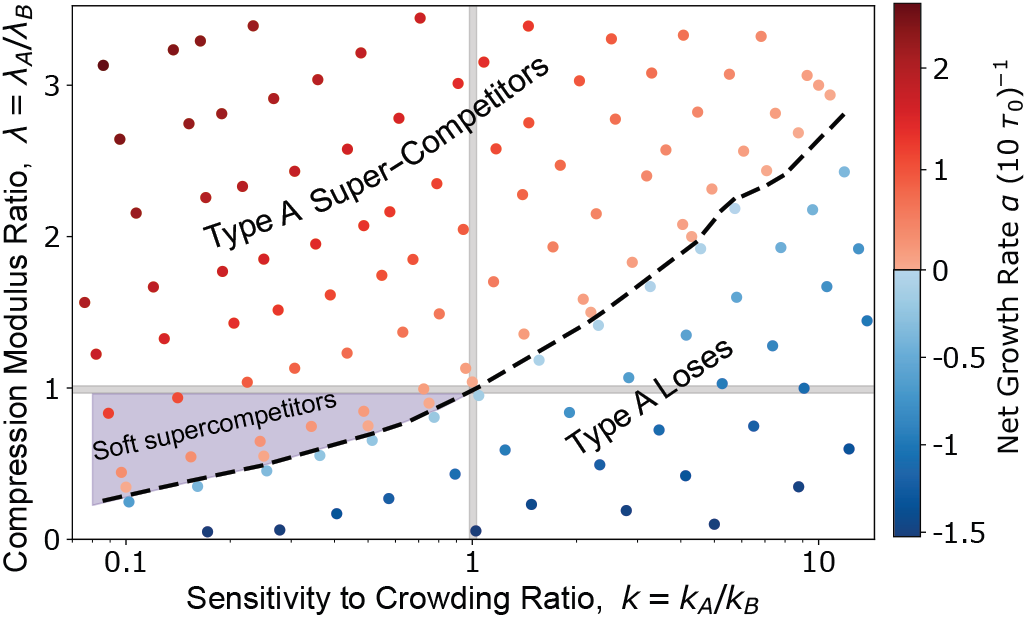
Trade-off between compressional elastic modulus and sensitivity to crowding determines outcome of cell competition. Phase diagram showing the net growth rate, *α*, of type A cells over the phase space of compressional elastic modulus ratio, *λ* = *λ*_*A*_*/λ*_*B*_, and the sensitivity to crowding ratio, *k* = *k*_*A*_*/k*_*B*_. The dashed black line represents where the net growth is approximately zero. Above this line type A is super-competitive, and below the line type A loses to type B. The purple region (*λ <* 1, *k <* 1, and above the dashed line) emphasizes where type A cells are more fit despite being softer than their type B neighbors. See tables I and II for a list of default model parameters.

### C. Mutant fitness is governed by physical parameters that regulate relative growth potential and susceptibility to apoptosis

Having identified the parameter regimes in *λ* -*k* phase space where the mutant type-A cells emerge as supercompetitors (Fig. **3**), we sought to investigate how perturbations in other physical cell parameters impact the fitness landscape. To understand how the fitness landscape transforms under parameter perturbations, we investigated seven representative type-A colonies (see Fig. **4**A) that traverse the border of super-competition and elimination. We then varied the model parameters that regulate cell proliferation and death dynamics for type A cells, including the cell growth rate, strength of homotypic adhesion relative to heterotypic adhesion, the G1 sizer threshold, sensitivities to density-dependant apoptosis and contact-dependant apoptosis.

**Fig. 4.**
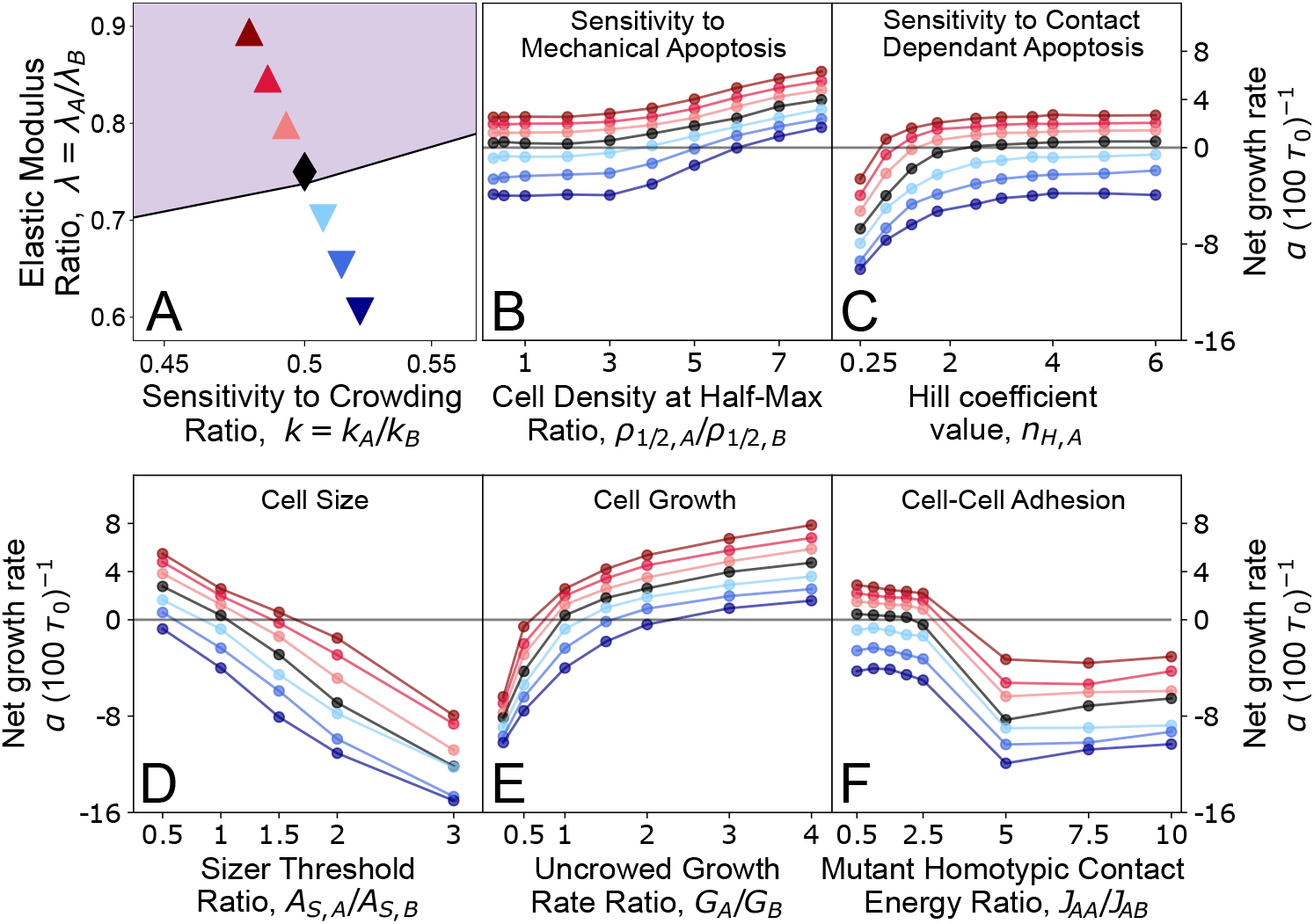
Mutant colony fitness is primarily determined by relative growth potential and susceptibility to apoptosis. (A) Points on the (*λ, k*) phase space, representing type A colonies with different values of elastic modulus ratio *λ*, and sensitivity to crowding ratio, *k*. Out of these seven representative cell colonies, 3 are supercompetitors (*α* > 0, purple shading), 3 are losers (*α* < 0, white shading), and one lies on the line of co-existence (black point). (B) Net growth rate of type-A colonies as a function of *ρ*_1*/*2,*A*_*/ρ*_1*/*2,*B*_, the ratio of the cell densities at half-maximum apoptosis probability (Eq. 5). Changing *ρ*_1*/*2,*A*_ alters the sensitivity to mechanical apoptosis of type A cells. (C) Net growth rate of type-A colonies as a function of *n*_*H,A*_, the hill coefficient in the equation for the probability of contact-dependent apoptosis (Eq. 6). Changing *n*_*H,A*_ alters the sensitivity to contact-dependent apoptosis of type A cells. (D) Net growth rate of type-A colonies as a function of the ratio of sizer threshold for type A and B cells. Cells that grow larger to progress through their cell cycle are less fit and more susceptible to apoptosis. (E) Net growth rate of type-A colonies as a function of the ratio of uncrowded growth rate for type A and B cells. Cells that grow faster are more fit and emerge as supercompetitors. (E) Net growth rate of type-A colonies as a function of the ratio of type A homotypic contact energy, *J*_*AA*_, over heterotypic contact energy of type A and B cells, *J*_*AB*_. For *J*_*AA*_*/J*_*AB*_ > 2.5, there is a substantial increase in heterotypic contact, leading to an increase in susceptibility of type A cells to contact-based apoptosis.

#### Sensitivity to mechanically-driven apoptosis

Differential sensitivity to mechanical pressure has been identified as a key factor in the elimination of cells through mechanical competition [8]. Cells with higher growth and proliferation rates exert pressure on their slow-growing neighbors, leading to an accumulation of mechanical stress [7]. If slower-growing cells are more sensitive to pressure or density-dependent apoptosis, they are more likely to be eliminated. For instance, in MDCK monolayers, *scribble* mutant cells are out-competed due to their heightened sensitivity to mechanical compaction [9, 25]. To test these behaviors in our model, we altered type A cell’s sensitivity to density-dependant apoptosis by varying the density *ρ*_1*/*2,*A*_ at half-maximum probability of apoptosis (Eq. 5). This means that if *ρ*_1*/*2,*A*_ *> ρ*_1*/*2,*B*_, then type A cells have a lower probability of death for any given value of local density. A sufficiently large difference in *ρ*_1*/*2_ values can effectively switch off density-dependent apoptosis for type A cells. We found substantial increases in the net growth rate of type A cells at higher values of the ratio *ρ*_1*/*2,*A*_*/ρ*_1*/*2,*B*_ (Fig. **4**B). For ratios *ρ*_1*/*2,*A*_*/ρ*_1*/*2,*B*_ ≥ 4, type A cells became supercompetitors, preferentially eliminating type B cells. In contrast, lower values of *ρ*_1*/*2,*A*_*/ρ*_1*/*2,*B*_, which effectively sets *P*_apo,den_ ≈ *p*_den,max_, had minimal effect on the net growth rate of type A cells.

#### Sensitivity to contact-based competition

Prior experiments on *myc*-driven competition in the *Drosophila* pupal notum have shown that cells expressing higher levels of *myc* preferentially eliminate wild-type cells when they are in mutual contact [12]. In quantitative terms, the probability of apoptosis for wild-type cells is proportional to the amount of heterotypic shared contact (Fig. **1**D-bottom). In our simulations, we altered the sensitivity to contact-dependant apoptosis by adjusting the Hill coefficient *n*_*H*_ in the probability equation (Eq. 6): 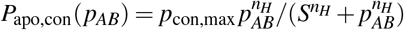 where the parameters *p*_con,max_ and *S* are obtained by fitting data [12] (Fig. **1**D-bottom). For a fixed value of *S*, increasing *n*_*H*_ would decrease the probability of apoptosis at a given value of the proportion of shared perimeter *p*_*AB*_. Thus, by changing the hill coefficient for type A cells, *n*_*H,A*_, the probability of apoptosis can be increased from its default value for *n*_*H,A*_ *<* 3.6 or decreased for *n*_*H,A*_ *>* 3.6 (Fig. **4**C). Interestingly, greatly decreasing the probability of apoptosis, by increasing *n*_*H,A*_, resulted in a negligible change in colony A fitness. This is consistent with our prior study which showed contact-dependant apoptosis dominated the outcome of competition only when the loser cells were fully mixed with the winner cells [31]. As expected, increasing the probability of contact-dependant apoptosis, by decreasing *n*_*H,A*_, gradually decreases the net growth rate of all tissues.

#### Perturbations to cell size and growth

Mutations in ribosomal proteins are one of the main drivers of cell competition [5, 51]. Reduced translation and slow growth affects cell size, suggesting that differential cell size and growth rate may be sensing cues for cell competition. We therefore asked how perturbations to cell size determines the emergence of supercompetitors. In our simulations, cell size is controlled by the G1/S sizer threshold, *A*_*S*_. The sizer threshold determines the minimum size necessary for a cell to progress from the G1 phase to the S/G2/M phase. In a crowded tissue, where growth is slow and dependant on the apoptosis of nearby cells to free space, the average cell cycle time is dominated by the duration of the G1 phase (Fig. **1**C). Alterations to cell size can tilt the balance between cell proliferation and apoptosis rates in two ways. First, a smaller *A*_*S*_ will reduce cell cycle times, thereby increasing proliferation rates. Secondly, larger cells will typically have more neighbors, hence a higher local density and a greater probability to die via mechanical and contact-based competition. We thus expect cell fitness to decrease with increasing cell size. In confirmation of this, we found an inverse linear relationship between the population net growth rate and sizer threshold for all type A colonies (Fig. **4**D). The increase in fitness for the smaller sizer threshold is primarily due to the shorter cell cycle times, representing an overall increase in proliferation rate relative to the rate of apoptosis. Consistent with this finding, *scribble* mutants that acquire a loser phenotype in competition with MDCK wild type cells, are typically larger in size [25, 31].

It has been suggested that differential growth rate can lead to the accumulation of mechanical stresses at the interface between two competing cell types [7, 23], leading to the elimination of the slower-growing cells via mechanically-driven apoptosis. To test this in our model, we varied the uncrowded growth rate *G* of type A cells (Eq. 3), the maximum growth rate in the absence of crowding. We found that altering the uncrowded growth rate of type A cells, *G*_*A*_, significantly impacted tissue fitness (Fig. **4**E), making type A cells either super-competitors for higher growth rates or losers for lower growth rates. From this result, in conjunction with Figure **3**, we find that the tissue growth potential, regulated by the parameters *λ, k*, and *G*, is crucial for determining the outcome of cell competition.

#### Cell-cell adhesion

Loss of cell-cell adhesions is a key feature of the epithelial-mesenchymal transition, during which epithelial cells acquire migratory and invasive characteristics [52]. While increased motility in supercompetitors can trigger apoptosis of suboptimal cells through contact-based elimination, weakened homotypic adhesions can also affect the mechanical properties of the mutant colonies, such as their compressibility. A recent study suggests that reduced cell-cell adhesions in HRas^V12^-expressing transformed cells contribute to their elimination by wild-type cells due to increased compressibility of the transformed cells [34].

To explore the role of intercellular adhesions in cell competition through our simulations, we varied the ratio of homotypic contact energy (*J*_*AA*_) to heterotypic contact energy (*J*_*AB*_), while keeping the other parameters fixed. Since a lower contact energy implies stronger adhesion, increasing *J*_*AA*_*/J*_*AB*_ weakens homotypic adhesion relative to heterotypic adhesion, thereby promoting greater intermixing between cell types A and B. We observed that for 0.5 *< J*_*AA*_*/J*_*AB*_ *<* 2.5, there is minimal effect on the net growth rate of colony A, despite incremental increases in heterotypic contact. However, when *J*_*AA*_*/J*_*AB*_ exceeds 2.5, a significant drop in net growth rate occurs as type A cells begin to repel one another, leading to type A cells migrating into the type B tissue and creating a fully mixed state (Video 4). In this state, heterotypic contact—and consequently, the probability of contact-dependent apoptosis—is maximized, resulting in the complete elimination of type A cells. Therefore, increasing homotypic contact energy (and thereby weakening homotypic adhesions) leads to the elimination of mutant cells at sufficiently high *J*_*AA*_*/J*_*AB*_ values.

### D. Critical size for colony growth and survival

The size of mutant colonies plays a crucial role in determining their growth, survival, and invasive potential. Previous theoretical studies have suggested a critical size threshold for the elimination of mutant colonies [15, 53–55], consistent with experiments on the influence of colony size on tumor invasion and cell removal [56, 57]. However, multiple factors contribute to whether a colony is eliminated by the surrounding tissue. Mechanistically, smaller colonies would experience a higher Laplace pressure (due to their higher interface curvature) compared to larger colonies, which may lead to their elimination through the compressive Laplace pressure [56]. This effect depends on the rigidity of the mutant colony compared to the surrounding tissue, as well as the homotypic contact energy that controls the colony surface tension. In addition, larger colonies, with their greater shared perimeter with the surrounding tissue, are more prone to elimination through contact-based competition. Therefore, mechanical and contact-based modes of cell elimination may either act antagonistically or act in synergy with each other depending on the geometry and mechanical properties of the tissue [58].

To understand how the interplay between colony size and mechanical properties impact their growth and survival, we varied the initial size of colony type A in our simulations across different values of the relative compressibility *λ* = *λ*_*A*_*/λ*_*B*_. We held the sensitivity to crowding ratio constant *k* = *k*_*A*_*/k*_*B*_ = 0.5, assuming that type A cells have a lower sensitivity to crowding, mimicking cancer cell colonies. The initial configuration consisted of a colony of type A cells of a specified radius (*R* = 50-300 pixels), positioned at the center and surrounded by type B cells (Fig. **5**A). We found that for a given value of relative compressibility *λ* (e.g., *λ* = 0.74 in Fig. **5**A), there is a critical colony size *R*∼ 125 pixels, below which the colony is eliminated, while above it, the colony continues to grow and survive (Videos 5-7). Here contact-based competition enabled type B tissue to eliminate the otherwise super-competitive type A colonies (for *λ* = 0.74, *k* = 0.5, see Fig. **3**), particularly when type A colonies were below a critical size. Thus, different initial sizes correspond to different sensitivities to contact-based elimination.

**Fig. 5.**
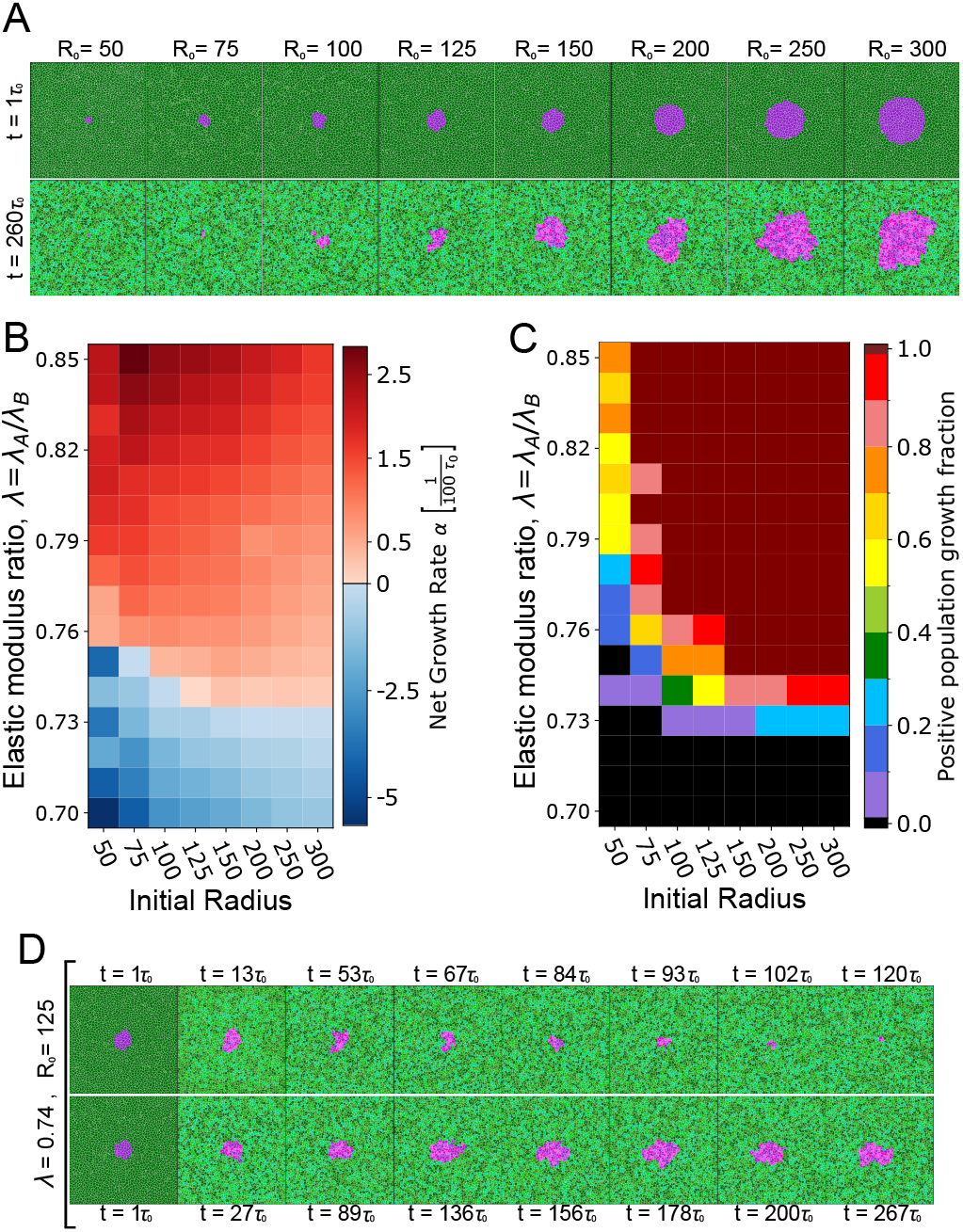
Critical colony size depends on relative cell compressibility. (A) Top row: Snapshots near the start of the simulation (*t* = *τ*_0_) depicting the initial size of colony A in terms of the radius (in pixels). Bottom row: Configuration of the system at *t* = 260*τ*_0_ showing either extinction or survival. (B) Colormap of net growth rate of colony A, as a function of different initial sizes and ratio of compressional moduli. The sensitivity to crowding ratio is set to *k* = 0.5, such that type A cells are less sensitive to crowding-induced suppression of growth. Each bin represents an average over 30 simulations. The net growth rate changes from negative to positive with increasing initial radius for *λ*_*A*_*/λ*_*B*_ in the range [0.74, 0.75]. (C) Phase diagram (colormap) showing the fraction of simulations which had positive population growth rate at *t* = 260*τ*_0_; that is *N*_*A*_(260*τ*_0_) *> N*_*A*_(0) where *N*_*A*_(*t*) is the population of the A colony at time *t*. (D) Time-evolution of a colony of initial size *R* = 125 pixels at *λ* = 0.74, showing an instance of elimination (top row) and survival (bottom row).

The critical colony size depends on the relative compressibility, *λ*, such that for *λ* = 0.75 cells exhibited a strictly positive net growth rate when the initial size *R >* 75 pixels (Fig.**5**B). For *λ*≤ 0.73, the average (*n* = 30) net growth rate is negative, implying that type A colonies would be eliminated regardless of their initial size (Fig.**5**B). In contrast, for *λ*≥0.76, the average net growth rate is positive, and type A colony invades by eliminating the type B cells. However, this behavior is probabilistic (Fig.**5**C), such that for *λ >* 0.73 there exists a compressibility-dependent critical size below which several simulations showed an increase in the population count of type A cells (Fig. **6**A). For the smallest colony size (*R* = 50 pixels, consisting of 6 type A cells), type B tissue managed to eliminate type A cells in some simulations, despite type A being more fit (*λ*≥ 0.76, Fig.**5**C and Fig. **6**B). Together, these results show that colony size impacts its fitness, but the critical colony size for growth is dependent on tissue mechanical properties and is governed by a probabilistic process.

**Fig. 6.**
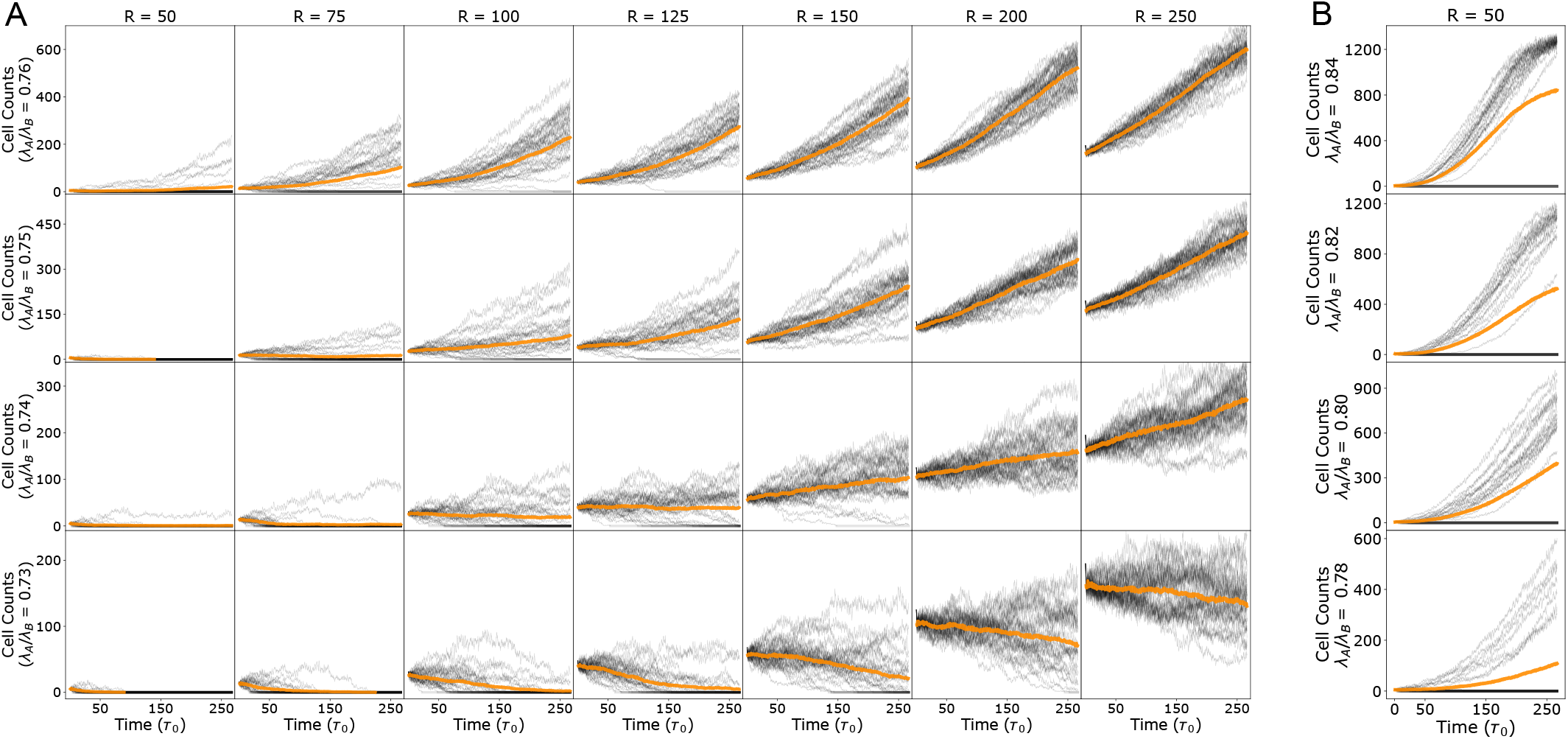
Population growth trajectories for varying initial sizes and relative compressibility. (A-B) Population growth trajectories (cell count vs. time) for Type-A colonies from individual simulations (black lines) and their average population (*n* = 30, orange lines). The parameters for Type-A colonies are indicated in each row with the relative compressibility *λ* = *λ*_*A*_*/λ*_*B*_ and by each column with the initial radius in pixel units. The sensitivity to crowding ratio is kept fixed at *k*_*A*_*/k*_*B*_ = 0.5. (A) Despite type A cells possessing a greater relative growth potential, for *λ* ≥ 0.74 and *k* = 0.5, smaller type A colonies can be outcompeted by the otherwise less fit type B tissue. This is evidenced by a decline in cell count in some simulations. For instance, note the comparisons: *λ* = 0.74 for *R* = 100 ⟶ 150, *λ* = 0.75 for *R* = 75 ⟶ 100, and *λ* = 0.76 for *R* = 50 ⟶ 75. (B) Population growth trajectories for initial size *R* = 50 pixels at different values of relative compressibility. Type B tissues can outcompete more fit type A colonies (see leftmost column of panel A), though with reduced frequency proportional to their relative compressibility.

The stochastic nature of cell colony elimination is illustrated in Fig.**5**D, where we see that two type-A cell colonies of the same initial size and identical physical properties show different outcomes in different simulation runs – one in which the colony survives (Fig.**5**D, bottom row) and the other where the colony is eliminated (Fig.**5**D, top row). The mechanism for elimination or invasion can be explained as follows (see Fig. **7**). When a cell at the boundary between two cell types dies, it permits a local invasion by the opposite type, with the extent of the invasion being slightly smaller than the typical cell size, 𝓁_0_. This invasion increases the proportion of heterotypic contact, particularly for the invading cells and the boundary cells adjacent to the invasion site. This increase in heterotypic contact would increase the probability of contact-dependent apoptosis, promoting further invasion. However, the invasion can be reversed if the opposite type’s tissue cells proliferate and replace the apoptotic boundary cells. Thus, a competition between proliferation and apoptosis of boundary cells determines whether the invasion will progress or be reversed.

**Fig. 7.**
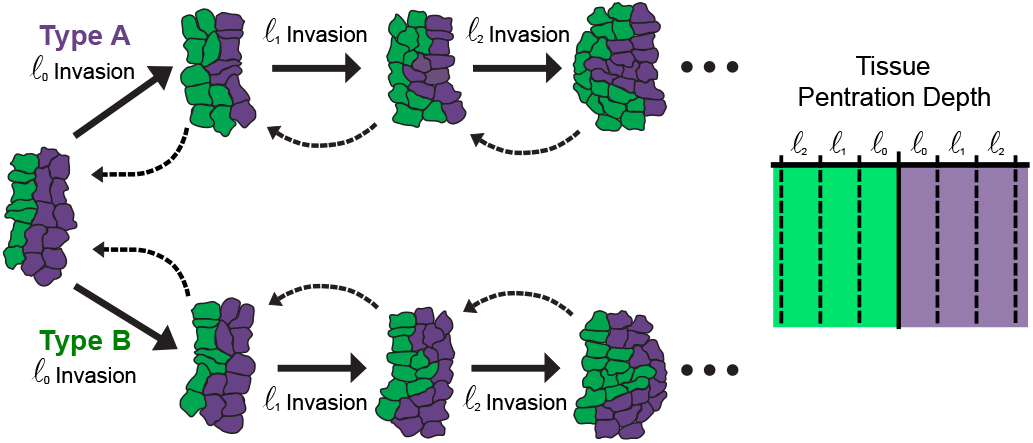
Colony invasion mechanism. Schematic showing the stochastic nature of invasion of a cell type onto another, initiated by a finger-like instability at the boundary between the two cell types. Invasions increase heterotypic contact yielding an average decrease in fitness for the small type A colonies due to a lack of replacement bulk cells. The death of a boundary cell allows for the initial invasion of the opposite type by some amount less than the typical cell size 𝓁_0_. This tends to increase the proportion of heterotypic shared contact, especially for each invading cell and the boundary cells neighboring the invasion. The invasion then progresses or is reversed in a probabilistic process that is most dependant on the ability of the tissue bulk cells to grow and replace the apoptotic boundary cells.

## DISCUSSION

In this study, we developed a cell-based model of cell competition that integrates cellular mechanics and decision-making rules using the framework of the cellular Potts model (Fig. **1**). The model allows for control over individual cell properties—such as mechanical forces, cell cycle parameters, and apoptosis rules—enabling us to identify the physical traits of individual cells that determine their potential to become supercompetitors. We explored this by embedding a mutant clone within a confluent tissue with distinct physical and biochemical properties, a situation relevant to developmental processes or early tumor formation.

Our findings demonstrate that differential compressibility alone can drive competition between otherwise identical cell types (Fig. **2**). Specifically, cells that are stiffer and more resistant to compression emerge as supercompetitors. This occurs because the stiffer cells compact the softer ones, increasing the local density of the softer cell type, making them more susceptible to elimination through density-dependent apoptosis. However, when cell types differ in their sensitivity to crowding or contact inhibition of proliferation, the outcome of competition is governed by a balance between compressibility and sensitivity to crowding. For example, increasing the sensitive to crowding parameter of the mutant cells generally leads to their elimination, even when they are stiffer than the wild-type cells. Interestingly, we identified a region in parameter space where softer mutant cells can outcompete wild-type cells that have a lower sensitivity to crowding (Fig. **3**). This finding is particularly relevant to cancer, as many cancerous cell lines are softer than their wild-type counterparts [59] and exhibit lower contact inhibition of proliferation [60]. Indeed, prior studies have shown that decreased cell stiffness correlates with increased malignancy in various cancers, including colon, ovarian, and breast cancers[46, 61–64].

The emergence of soft supercompetitors with low crowding sensitivity can also be influenced by other single-cell parameters, such as those regulating proliferative fitness or susceptibility to apoptosis (Fig. **4**). For example, reducing the sensitivity to density-dependent or contact-dependent apoptosis can promote supercompetitive behavior in parameter regimes where mutant colonies would otherwise be driven to extinction. Similarly, enhancing proliferative fitness by increasing growth rates or reducing the G1 size threshold can also induce supercompetitive behavior. Another key physical factor is the ratio of homotypic to heterotypic contact energy (*J*_*AA*_*/J*_*AB*_), which reflects the relative adhesion between cell types A and B compared to adhesion within type A cells. Higher *J*_*AA*_*/J*_*AB*_ values promote greater intermixing of cell types, increasing heterotypic contact and thus making type A cells more vulnerable to elimination through contact-dependent apoptosis. This aligns with experimental evidence showing that increased cohesion within loser cells protects them from elimination, whereas greater intermixing between winner and loser cells leads to enhanced elimination of the losers [12].

The initial size of a mutant colony plays a crucial role in its survival probability, depending on its compressibility (Fig. **5**). If the mutant cells are softer, colony growth is inhibited regardless of initial size. However, when compressibility exceeds a critical threshold, there is a critical initial size above which the colony is more likely to invade. Below this critical size, colony survival is stochastic, driven by a finger-like instability at the boundary between cell types (Fig. **7**). Local apoptosis in the mutant clone creates space for wild-type cell proliferation, promoting invasion into the mutant colony. This invasion increases heterotypic contact, which in turn drives further apoptosis and invasion. Interestingly, our model predicts that this invasion can be reversed if the mutant clone has a higher compressional modulus or lower contact inhibition of proliferation, highlighting the importance of physical cell parameters in determining supercompetitive behavior.

## ACKNOWLEDGEMENTS

We thank Fernanda Perez-Verdugo, Daniel Gradeci and Guillaume Charras for useful discussions. SB acknowledges support from the National Institutes of Health (NIH R35 GM143042).

## APPENDIX A: CELLULAR POTTS MODEL SIMULATION

We utilized the Cellular Potts Model (CPM) [38] to simulate the dynamics of cell competition. The CPM is a versatile computational framework widely used to model biological processes such as cell growth, migration, and tissue morphogenesis. The model operates by running a Monte Carlo simulation using the Metropolis-Hastings algorithm [40, 41], which minimizes the system’s free energy, as described in the main text. While alternative cell-based modeling approaches exist, including vertex models, cell-center models, and particle-based models, we selected the CPM for its simplicity in integrating key components of this study, such as agent-based rules, boundary conditions, and cell-extracellular medium interactions. The simulations were conducted using CompuCell3D [39], a software package that implements the CPM framework and the Metropolis-Hastings algorithm. The version of CompuCell3D used in this study allows for cell fragmentation based on lattice temperature, which we set to *T* = 10 in lattice energy units. This temperature was found to result in negligible fragmentation. Simulations are initiated from confluent type B tissue in homeostasis on a 2D lattice of 1500×1500 pixels. At time *t* = 0, cells within a radius of 300 pixels (default value for initial radius) of the center of the tissue are transformed to type A cells (initial population counts are *N*_*A*_ = 284 cells and *N*_*B*_ = 1931 cells). Other default parameters are specified in Tables I and II. Custom simulation codes are available at https://github.com/BanerjeeLab/CellCompetition.

## APPENDIX B: RULES FOR CELL ELIMINATION

In our simulations, cell elimination involves two mechanisms: programmed cell death (apoptosis) and live-cell extrusion. The specific mathematical rules for these processes are described in the main text, Section II.B.

In our simulations, the probability of apoptosis through mechanical and contact-based competition is calculated every 10 Monte Carlo steps, which, on the time scale of the timer threshold, reproduces the experimentally observed probabilities. When a cell is eliminated through apoptosis, its target area is set to zero, causing it to shrink rapidly. During this process, surrounding cells grow to occupy the space left by the apoptotic cell. Within the simulation, this is implemented as:

1. Iterate over each cell
2. Calculate the local density for the given cell, *ρ*_*c*_ (eq 4) and determine the shared perimeter with cells of a different type.
3. Check whether the cell undergoes mechanical apoptosis with probability *P*(*ρ*_*c*_) (Eq. 5) or contact-based apoptosis (Eq. 6).
4. If the cell is apoptotic, set the target area and elastic modulus to *A*_*T*_ = 0 and *λ* = 5, which would promote rapid, but not instantaneous, shrinkage.

In addition to apoptosis, cells may be eliminated via live-cell extrusions. This process occurs when a cell’s area decreases significantly relative to the average population area. Experimental observations [18] indicate that extrusion is triggered by compressive stress within the tissue, causing the cell to detach from the substrate and be ejected from the monolayer. In our simulations, any cell whose area falls below ⟨ *A*⟩ */*4 (where ⟨*A*⟩ is the average area of the colony) is immediately eliminated, mimicking the rapid time scale of extrusion [18, 65]. This is implemented in the simulation as follows:

1. Calculate the average cellular area, ⟨*A*⟩, within the tissue.
2. Iterate over each cell and check whether its area, *A*, satisfies *A <* ⟨*A*⟩ */*4. Any cell meeting this criterion is immediately ejected by converting its lattice sites into medium sites.

## APPENDIX C: MODEL PARAMETERIZATION

To compare our simulations with physiological scenarios, converting lattice units in the CPM framework to physical units is essential. This involves two primary conversions: translating pixels to SI units for distance, and Monte Carlo Steps (mcs) to SI units for time. To determine the distance conversion, we compared the experimentally reported average cell volume for MDCK cells during subconfluence [31, 43, 66] with the average subconfluent colony area in our simulations, *Ā*_*SC*_ = 3200 pixels. This yielded a conversion factor: 1 pixel = (0.13 ± 0.03)*µm*^2^. Applying this factor to the G1/S size threshold gave us *A*_*S*_ = 1377 pixels = (179.01± 41.31)*µm*^2^. Based on this, we set the default sizer threshold to *A*_*S*_ = 1400 pixels. We normalized simulation time with the average cell cycle duration in uncrowded conditions, i.e., the cell’s timer duration *τ*_0_. Experimental data for typical epithelial cells suggest that the S/G2/M phase lasts about *τ*_0_ ≈ 10 hours [43, 66]. This provides the timescale conversion from Monte Carlo steps to hours, *τ*_0_ = 225 mcs ≈ 10 hr (or 1 mcs ≈ 2.67 min).

The single-cell growth rate, *G*, is sampled from a Gaussian distribution and determines the change in target area per Monte Carlo step (Eq. (3)). The mean value, *G*_0_, is set so that isolated cells maintain the average subconfluent area *ĀSC*, with 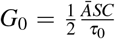. The standard deviation is set to 5% *G*_0_. While *G* controls the growth rate (d*A*^*T*^ */*d*t*) for isolated cells, crowding in the tissue restricts available space, causing the actual cell area to deviate from the target. This activates the exponential term in Eq. (3),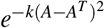, driving d*A*^*T*^ */*d*t* to zero until the cell area increases. The parameter *k*, representing the sensitivity to crowding or contact inhibitio of proliferation, quantifies the allowed deviation between the actual and target areas. The default value, *k* = 0.01 pixels^-2^, was empirically chosen to ensure appropriate colony growth dynamics over several cell cycles.

In our CPM model, five parameters define the energy scale in the Hamiltonian: the elastic energy term *λ*, and the contact energy terms *J*_AA_, *J*_BB_, *J*_AB_, and *J*_AM_ = *J*_BM_ = *J*_MM_. Here, *λ* represents the area elasticity modulus, set to a default value of *λ* = 1. The *J* parameters represent the strength of intercellular adhesions; higher values of *J* indicate lower adhesion and higher surface tension. Accordingly, we set *J*_AA_ = 12.5, *J*_AB_ = 10, *J*_BB_ = 6, and *J*_AM_ = *J*_BM_ = *J*_MM_ = 15, allowing cells to adhere more strongly to each other than to the medium. The variations in *J*_AA_, *J*_BB_, and *J*_AB_ resulted in different morphologies for the growing colony of type A cells.

## APPENDIX D: DESCRIPTION OF VIDEOS

### Video Legends

Each video depicts a simulation of a type A colony embedded in a type B tissue. The cells change color upon division to make these events easier to see. Both cell types cycle through four colors:

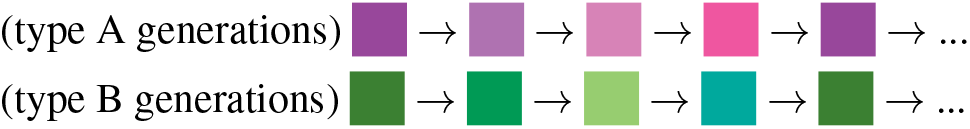

**Video 1**. This video shows both cell types with the default parameter values. That is, with the exception cell-cell adhesion, they are mechanically identical. This results in the type A colony maintaining approximately the same size.

**Video 2**. This video shows that increasing the stiffness of type A (*λ*_*A*_*/λ*_*B*_ = 2), results in the type A colony rapidly outgrowing the type B tissue.

**Video 3**. This video shows that decreasing the stiffness of type A (*λ*_*A*_*/λ*_*B*_ = 0.5), results in the elimination of the type A colony.

**Video 4**. This video shows an example of the type A colony dispersing to a mixed state and eventually eliminated due to repulsion induced by setting *J*_*AA*_ = 40.

**Video 5**. This video shows an example of the type A colony, with an initial radius *R* = 100 pixels and parameters (*λ*_*A*_*/λ*_*B*_, *k*_*A*_*/k*_*B*_) = (0.74, 0.5), being out competed over a long time span (161*τ*_0_).

**Video 6**. This video shows an example of the type A colony, with an initial radius *R* = 125 pixels and parameters (*λ*_*A*_*/λ*_*B*_, *k*_*A*_*/k*_*B*_) = (0.74, 0.5), deforming in shape while maintaining the size over a long time span (266*τ*_0_).

**Video 7**. This video shows an example of the type A colony, with an initial radius *R* = 200 pixels and parameters (*λ*_*A*_*/λ*_*B*_, *k*_*A*_*/k*_*B*_) = (0.74, 0.5), deforming in shape and growing in size over a long time span (266*τ*_0_).

## References

[1] R. Levayer and E. Moreno, Mechanisms of cell competition: themes and variations, Journal of Cell Biology 200, 689 (2013).

[2] J.-P. Vincent, A. G. Fletcher, and L. A. Baena-Lopez, Mechanisms and mechanics of cell competition in epithelia, Nature Reviews Molecular Cell Biology 14, 581 (2013).

[3] M. M. Merino, R. Levayer, and E. Moreno, Survival of the fittest: essential roles of cell competition in development, aging, and cancer, Trends in cell biology 26, 776 (2016).

[4] E. Moreno, Is cell competition relevant to cancer?, Nature Reviews Cancer 8, 141 (2008).

[5] G. Morata and P. Ripoll, Minutes: Mutants of drosophila autonomously affecting cell division rate, Developmental Biology 42, 211 (1975).

[6] E. Moreno and K. Basler, endmyc transforms cells into super-competitors, Cell 117, 117 (2004).

[7] B. I. Shraiman, Mechanical feedback as a possible regulator of tissue growth, Proceedings of the National Academy of Sciences 102, 3318 (2005).

[8] A. Matamoro-Vidal and R. Levayer, Multiple influences of mechanical forces on cell competition, Current Biology 29, R762 (2019).

[9] L. Wagstaff, M. Goschorska, K. Kozyrska, G. Duclos, I. Kucinski, A. Chessel, L. Hampton-O’Neil, C. R. Bradshaw, G. E. Allen, E. L. Rawlins, P. Silberzan, R. E. Carazo Salas, and Piddini, Mechanical cell competition kills cells via induction of lethal p53 levels, Nature Communications 7, 11373 (2016).

[10] R. Levayer, C. Dupont, and E. Moreno, Tissue crowding induces caspase-dependent competition for space, Current Biology 26, 670 (2016).

[11] E. Moreno, K. Basler, and G. Morata, Cells compete for decapentaplegic survival factor to prevent apoptosis in drosophila wing development, Nature 416, 755 (2002).

[12] R. Levayer, B. Hauert, and E. Moreno, Cell mixing induced by myc is required for competitive tissue invasion and destruction, Nature 524, 476 (2015).

[13] C. Díaz-Díaz, L. F. de Manuel, D. Jimenez-Carretero, M. C. Montoya, C. Clavería, and M. Torres, Pluripotency surveillance by myc-driven competitive elimination of differentiating cells, Developmental Cell 42, 585 (2017).

[14] M. Yamamoto, S. Ohsawa, K. Kunimasa, and T. Igaki, The ligand sas and its receptor ptp10d drive tumour-suppressive cell competition, Nature 542, 246 (2017).

[15] M. Basan, T. Risler, J. Joanny, X. Sastre-Garau, and J. Prost, Homeostatic competition drives tumor growth and metastasis nucleation, HFSP Journal 3, 265 (2009).

[16] M. Abercrombie, Contact inhibition and malignancy, Nature 281, 259 (1979).

[17] A. Puliafito, L. Hufnagel, P. Neveu, S. Streichan, A. Sigal, D. K. Fygenson, and B. I. Shraiman, Collective and single cell behavior in epithelial contact inhibition, Proceedings of the National Academy of Sciences 109, 739 (2012).

[18] G. T. Eisenhoffer, P. D. Loftus, M. Yoshigi, H. Otsuna, C.-B. Chien, P. A. Morcos, and J. Rosenblatt, Crowding induces live cell extrusion to maintain homeostatic cell numbers in epithelia, Nature 484, 546 (2012).

[19] S. J. Streichan, C. R. Hoerner, T. Schneidt, D. Holzer, and L. Hufnagel, Spatial constraints control cell proliferation in tissues, Proceedings of the National Academy of Sciences 111, 5586 (2014).

[20] K. D. Irvine and B. I. Shraiman, Mechanical control of growth: ideas, facts and challenges, Development 144, 4238 (2017).

[21] I. Di Meglio, A. Trushko, P. Guillamat, C. Blanch-Mercader, S. Abuhattum, and A. Roux, Pressure and curvature control of the cell cycle in epithelia growing under spherical confinement, Cell Reports 40 (2022).

[22] B. Zhao, X. Wei, W. Li, R. S. Udan, Q. Yang, J. Kim, J. Xie, T. Ikenoue, J. Yu, L. Li, P. Zheng, K. Ye, A. Chinnaiyan, G. Halder, Z.-C. Lai, and K.-L. Guan, enInactivation of YAP oncoprotein by the hippo pathway is involved in cell contact inhibition and tissue growth control, Genes Dev 21, 2747 (2007).

[23] Y. Pan, I. Heemskerk, C. Ibar, B. I. Shraiman, and K. D. Irvine, Differential growth triggers mechanical feedback that elevates hippo signaling, Proceedings of the National Academy of Sciences 113, E6974 (2016).

[24] N.-G. Kim, E. Koh, X. Chen, and B. M. Gumbiner, E-cadherin mediates contact inhibition of proliferation through hippo signaling-pathway components, Proceedings of the National Academy of Sciences 108, 11930 (2011).

[25] A. Bove, D. Gradeci, Y. Fujita, S. Banerjee, G. T. Charras, and A. R. Lowe, Local cellular neighborhood controls proliferation in cell competition, Molecular Biology of the Cell 28, 3215 (2017).

[26] D. Gradeci, A. Bove, G. Charras, A. R. Lowe, and S. Banerjee, Single-cell approaches to cell competition: high-throughput imaging, machine learning and simulations, in Seminars in Cancer Biology, Vol. 63 (Elsevier, 2020) pp. 60–68.

[27] S. Nishikawa, A. Takamatsu, S. Ohsawa, and T. Igaki, Mathematical model for cell competition: Predator–prey interactions at the interface between two groups of cells in monolayer tissue, Journal of Theoretical Biology 404, 40 (2016).

[28] S. Nishikawa and A. Takamatsu, Effects of cell death-induced proliferation on a cell competition system, Mathematical Biosciences 316, 108241 (2019).

[29] R. J. Murphy, P. R. Buenzli, R. E. Baker, and M. J. Simpson, Mechanical cell competition in heterogeneous epithelial tissues, Bulletin of Mathematical Biology 82, 130 (2020).

[30] A. Tsuboi, S. Ohsawa, D. Umetsu, Y. Sando, E. Kuranaga, T. Igaki, and K. Fujimoto, Competition for space is controlled by apoptosis-induced change of local epithelial topology, Current Biology 28, 2115 (2018).

[31] D. Gradeci, A. Bove, G. Vallardi, A. R. Lowe, S. Banerjee, and G. Charras, Cell-scale biophysical determinants of cell competition in epithelia, eLife 10, e61011 (2021).

[32] T. F. Pak, J. Pitt-Francis, and R. E. Baker, A mathematical framework for the emergence of winners and losers in cell competition, Journal of Theoretical Biology 577, 111666 (2024).

[33] X. Li, A. Datta, and S. Banerjee, Proliferation symmetry breaking in growing tissues, bioRxiv 10.1101/2024.09.03.610990 (2024).

[34] P. Gupta, S. Kayal, N. Tanimura, S. P. Pothapragada, H. K. Senapati, P. Devendran, Y. Fujita, D. Bi, and T. Das, Mechanical imbalance between normal and transformed cells drives epithelial homeostasis through cell competition 10.1101/2023.09.27.559723 (2023).

[35] F. Bosveld, B. Guirao, Z. Wang, M. Rivière, I. Bonnet, F. Graner, and Y. Bellaïche, Modulation of junction tension by tumor-suppressors and proto-oncogenes regulates cell-cell contacts, Development 10.1242/dev.127993 (2016).

[36] S.-W. Lee and Y. Morishita, Two types of critical cell density for mechanical elimination of abnormal cell clusters from epithelial tissue, PLOS Computational Biology 18, e1010178 (2022).

[37] L. C. Carpenter, F. Pérez-Verdugo, and S. Banerjee, Mechanical control of cell proliferation patterns in growing epithelial monolayers, Biophysical Journal 123, 909–919 (2024).

[38] F. m. c. Graner and J. A. Glazier, Simulation of biological cell sorting using a two-dimensional extended potts model, Phys. Rev. Lett. 69, 2013 (1992).

[39] M. H. Swat, G. L. Thomas, J. M. Belmonte, A. Shirinifard, D. Hmeljak, and J. A. Glazier, Multi-scale modeling of tissues using compucell3d, in Methods in Cell Biology, Vol. 110 (Elsevier, 2012) pp. 325–366.

[40] N. Metropolis, A. W. Rosenbluth, M. N. Rosenbluth, A. H. Teller, and E. Teller, Equation of State Calculations by Fast Computing Machines, The Journal of Chemical Physics 21, 1087 (2004).

[41] W. K. Hastings, Monte Carlo sampling methods using Markov chains and their applications, Biometrika 57, 97 (1970).

[42] S. Xie and J. M. Skotheim, A g1 sizer coordinates growth and division in the mouse epidermis, Current Biology 30, 916 (2020).

[43] J. Devany, M. J. Falk, L. J. Holt, A. Murugan, and M. L. Gardel, Epithelial tissue confinement inhibits cell growth and leads to volume-reducing divisions, Developmental Cell (2023).

[44] E. E. Bastounis, F. Serrano-Alcalde, P. Radhakrishnan, P. Engström, M. J. Gómez-Benito, M. S. Oswald, Y.-T. Yeh, J. G. Smith, M. D. Welch, J. M. García-Aznar, et al., Mechanical competition triggered by innate immune signaling drives the collective extrusion of bacterially infected epithelial cells, Developmental Cell 56, 443 (2021).

[45] L. Wullkopf, A.-K. V. West, N. Leijnse, T. R. Cox, C. D. Madsen, L. B. Oddershede, and J. T. Erler, Cancer cells’ ability to mechanically adjust to extracellular matrix stiffness correlates with their invasive potential, Molecular Biology of the Cell 29, 2378 (2018).

[46] W. Xu, R. Mezencev, B. Kim, L. Wang, J. McDonald, and T. Sulchek, Cell stiffness is a biomarker of the metastatic potential of ovarian cancer cells, PLOS ONE 7, 1 (2012).

[47] H.-H. Lin, H.-K. Lin, I.-H. Lin, Y.-W. Chiou, H.-W. Chen, C.-Y. Liu, I. Hans, C. Harn, W.-T. Chiu, Y.-K. Wang, et al., Mechanical phenotype of cancer cells: cell softening and loss of stiffness sensing, Oncotarget 6, 20946 (2015).

[48] T. Fuhs, F. Wetzel, A. W. Fritsch, X. Li, R. Stange, S. Pawlizak, T. R. Kießling, E. Morawetz, S. Grosser, F. Sauer, et al., Rigid tumours contain soft cancer cells, Nature Physics 18, 1510 (2022).

[49] B. St. Croix, C. Sheehan, J. W. Rak, V. A. Flørenes, J. M. Slingerland, and R. S. Kerbel, E-cadherin–dependent growth suppression is mediated by the cyclin-dependent kinase inhibitor p27kip1, The Journal of cell biology 142, 557 (1998).

[50] A. I. McClatchey and A. S. Yap, enContact inhibition (of proliferation) redux, Curr Opin Cell Biol 24, 685 (2012).

[51] M. Kiparaki and N. E. Baker, Ribosomal protein mutations and cell competition: autonomous and nonautonomous effects on a stress response, Genetics 224, iyad080 (2023).

[52] M. Janiszewska, M. C. Primi, and T. Izard, Cell adhesion in cancer: Beyond the migration of single cells, Journal of Biological Chemistry 295, 2495 (2020).

[53] H. B. Frieboes, M. E. Edgerton, J. P. Fruehauf, F. R. Rose, L. K. Worrall, R. A. Gatenby, M. Ferrari, and V. Cristini, Prediction of drug response in breast cancer using integrative experimental/computational modeling, Cancer Research 69, 4484 (2009).

[54] D. Drasdo and S. Höhme, A single-cell-based model of tumor growth in vitro: monolayers and spheroids, Physical Biology 2, 133 (2005).

[55] J. Li, S. K. Schnyder, M. S. Turner, and R. Yamamoto, Competition between cell types under cell cycle regulation with apoptosis, Physical Review Research 4, 033156 (2022).

[56] C. Bielmeier, S. Alt, V. Weichselberger, M. La Fortezza, H. Harz, F. Jülicher, G. Salbreux, and A.-K. Classen, Interface contractility between differently fated cells drives cell elimination and cyst formation, Current Biology 26, 563 (2016).

[57] B. D. Riehl, E. Kim, T. Bouzid, and J. Y. Lim, The role of microenvironmental cues and mechanical loading milieus in breast cancer cell progression and metastasis, Frontiers in Bio-engineering and Biotechnology 8, 608526 (2021).

[58] R. Levayer, Solid stress, competition for space and cancer: The opposing roles of mechanical cell competition in tumour initiation and growth, in Seminars in Cancer Biology, Vol. 63 (Elsevier, 2020) pp. 69–80.

[59] C. Alibert, B. Goud, and J.-B. Manneville, enAre cancer cells really softer than normal cells?, Biol. Cell 109, 167 (2017).

[60] D. Hanahan and R. Weinberg, Hallmarks of cancer: The next generation, Cell 144, 646 (2011).

[61] M. Pachenari, S. M. Seyedpour, M. Janmaleki, S. Babazadeh Shayan, S. Taranejoo, and H. Hosseinkhani, enMechanical properties of cancer cytoskeleton depend on actin filaments to microtubules content: investigating different grades of colon cancer cell lines, J. Biomech. 47, 373 (2014).

[62] V. Swaminathan, K. Mythreye, E. T. O’Brien, A. Berchuck, G. C. Blobe, and R. Superfine, enMechanical stiffness grades metastatic potential in patient tumor cells and in cancer cell lines, Cancer Res. 71, 5075 (2011).

[63] J. Guck, S. Schinkinger, B. Lincoln, F. Wottawah, S. Ebert, M. Romeyke, D. Lenz, H. M. Erickson, R. Ananthakrishnan, D. Mitchell, J. Käs, S. Ulvick, and C. Bilby, enOptical deformability as an inherent cell marker for testing malignant transformation and metastatic competence, Biophys. J. 88, 3689 (2005).

[64] N. Gal and D. Weihs, enIntracellular mechanics and activity of breast cancer cells correlate with metastatic potential, Cell Biochem. Biophys. 63, 199 (2012).

[65] L. Kocgozlu, T. Saw, A. Le, I. Yow, M. Shagirov, E. Wong, R.-M. Mège, C. Lim, Y. Toyama, and B. Ladoux, Epithelial cell packing induces distinct modes of cell extrusions, Current Biology 26, 2942 (2016).

[66] C. Cadart, S. Monnier, J. Grilli, P. J. Sáez, N. Srivastava, R. Attia, E. Terriac, B. Baum, M. Cosentino-Lagomarsino, and M. Piel, Size control in mammalian cells involves modulation of both growth rate and cell cycle duration, Nature communications 9, 3275 (2018).

